# Multiscale modelling of motility wave propagation in cell migration

**DOI:** 10.1101/2020.01.28.924191

**Authors:** Hamid Khatee, Andras Czirok, Zoltan Neufeld

**Affiliations:** School of Mathematics and Physics, The University of Queensland, St. Lucia, Brisbane, QLD 4072, Australia; Department of Biological Physics, Eotvos University, Budapest, 1053, Hungary; Department of Anatomy and Cell Biology, University of Kansas Medical Center, Kansas City, KS 66160, USA

## Abstract

The collective motion of cell monolayers within a tissue is a fundamental biological process that occurs during tissue formation, wound healing, cancerous invasion, and viral infection. Experiments have shown that at the onset of migration, the motility is self-generated as a polarization wave starting from the leading edge of the monolayer and progressively propagates into the bulk. However, it is unclear how the propagation of this motility wave is influenced by cellular properties. Here, we investigate this using a computational model based on the Potts model coupled to the dynamics of intracellular polarization. The model captures the propagation of the polarization wave initiated at the leading edge and suggests that the cells cortex can regulate the migration modes: strongly contractile cells may depolarize the monolayer, whereas less contractile cells can form swirling movement. Cortical contractility is further found to limit the cells motility, which (i) decelerates the wave speed and the leading edge progression, and (ii) destabilises the leading edge into migration fingers. Together, our model describes how different cellular properties can contribute to the regulation of collective cell migration.

## 1 Introduction

Collective cell migration is a fundamental process both during embryonic development and pathophysiology such as wound healing or cancer metastasis^1–3^. Conceptually, the dynamics of the migrating cells have been best understood in epithelial monolayers^4,5^ which is the simplest tissue that line organs throughout the body^6^. As cells cooperatively move together^7,8^, each cell needs to establish its own polarity: confining certain biochemical processes to the front of the cell while others to the rear^9–11^. The most fundamental hallmark of cell polarization manifests in cytoskeletal dynamics: polymerization of F-actin at the leading edge, a process coordinated by a variety of intracellular signaling molecules^10^. Actin polymerization and the activity of myosin II molecular motors produces cytoskeletal flows which translate to cell motility^1,7,12–16^.

The synchronization of cell polarity during collective migration as well as the guidance from mechanical environmental cues are poorly understood^17^. The process likely involves cell-cell junctions are mediated by adhesion proteins (e.g., transmembrane receptor E-cadherin) coupled to the contractile actomyosin cytoskeleton through cytoplasmic adaptor proteins (e.g., *α*-catenin)^4^. Actomyosin contractility at the cell cortex can modulate both adhesion strength^18^, and cell shape, both exerting an important influence on tissue remodelling^1,6^.

A common experimental system to study collective cell migration *in vitro* is the wound closure assay, in which a barrier, the "wound", divides a monolayer culture. After removal of the barrier, cells migrate into the empty region and eventually create or restore a continuous monolayer of cells. Live imaging of such assays^19–21^ indicated that the number of cells in the cell-free region increases, mainly due to active cell migration and not proliferation^19–22^. Furthermore, the onset of migration is delayed for the cells deep in the bulk compared to those in the vicinity of the front boundary. In several cultures the onset of motility can be observed as a polarity wave propagating backwards from the edge of epithelial monolayer^19–21^.

Several theoretical models have been proposed to explain the coordination between cells during collective migration^20,22–34^. However, it remains elusive how the inter- and intra-cellular mechanobiology regulates the initiation and propagation of the polarization wave through a monolayer of cells. Recently, we developed a one-dimensional model of this mechanism, which involved mechanical forces and biomechanical feedback between cells. The model predicted a traveling wave that transmits polarization information and initiates motility in the bulk of the monolayer^35^. Here we extend our model to two dimensions and computationally study the effects of cellular mechanical properties on the expansion of a confluent epithelial monolayer. We develop a multiscale model for active cell movement, without cell proliferation, to predict collective motility in barrier-removal assays. We show that the model reproduces experimentally observed modes of migratory dynamics, and provides experimentally testable predictions how various mechanical properties modulate the collective migration of cells.

## 2 The model

A barrier-removal assay is often utilized to study collective motility of epithelial cells. Our model represents the migration of cells toward the cell-free region, after the barrier removal – and focuses on the role of intercellular interactions and intracellular mechanics in the process. The cell-cell interactions are represented using the Cellular Potts Model (CPM)^23,36^. The CPM is a lattice model which is computationally and conceptionally simpler than most off-lattice models (e.g., vertex model), while it provides a realistic description of cell shapes^37,38^. The intra-cellular polarity dynamics was formulated as a set of ODEs, coupled to each model cell of the CPM. This representation of cell motility is a generalisation of the model we used in the context of a simple one-dimensional chain of cells for the onset of collective cell movement^35^. The model is implemented using the open-source software package CompuCell3D (CC3D)^39^.

### 2.1 Inter-cellular dynamics

The CPM represents the cells on a two-dimensional lattice, where each cell covers a set of connected lattice sites or pixels; each pixel can only be occupied by one cell at a time. In this paper, the lattice is a rectangular surface (1500 and 320 pixels in the *x*- and *y*-dimensions, respectively). The expansion and retraction of the cells boundaries is determined by minimizing a phenomenological energy or goal-function *E*, defined in terms of the area *A*_*σ*_ and perimeter *L*_*σ*_ of *N* cells (indices *σ* = 1, …, *N*)^33,40^, and the motile forces 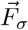 modeled as:

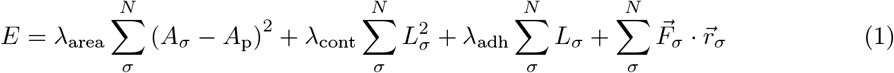

The first term of Equation (1) models the compressibility of cell body by penalizing the deviation of cell areas from a pre-set value *A*_p_, 100 pixels (see Table 1). The second term represents the contractility of the cell cortex (perimeter) *L*_*σ*_ as a spring with zero equilibrium length. The penalty parameter *λ*_cont_ represents cortical actomyosin contractility, in the vicinity of lateral cell membranes^41^. The third term describes the cell-cell adhesion mediated by adhesion molecules, such as E-cadherin. Note that *λ*_adh_ < 0 to represent that cells preferentially expand their boundaries shared with neighbouring cells. This is however balanced by the contractile tension along the cell cortex. The last term represents a motile force driving cells into the direction of 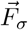, where 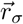 is the position vector of the cells center of mass. The prefactors *λ*_area_, *λ*_cont_, *λ*_adh_, and *F*_max_ reflect the relative importance of the corresponding cellular properties to set cellular morphology.

**Table 1:** Model parameters.

| Parameter | Value |
| --- | --- |
| Number of cells, $N$ | $3000^{\ddagger}$ |
| Cell size (pixel × pixel) | $10 \times 10$ |
| Initial cell area, $A_{\sigma}$ (pixel × pixel) | 100 |
| Preferred area, $A_p$ (pixel × pixel) | 100 |
| Area strain, $\lambda_{\text{area}}$ | 70 |
| Temperature, $T$ | 50 |
| Depolarization rate, $\beta$ | 0.1 |
| Polarization rate, $\gamma$ | 1 |
| Half-saturation constant, $\alpha$ | 1 |
| Hill coefficient, $n$ | 10 |
| Maximal motile force, $F_{\max}$ | 1000 |
| Initial polarity magnitude of leading cells, $p_0$ | 4 |

‡ 100 and 30 cells in the $x$ - and $y$ -dimensions, respectively.

The dynamics of the CPM is defined by a stochastic series of elementary steps, where a cell expands or shrinks accommodated by a corresponding area change in the adjacent cell (or empty area)^36,39^. The algorithm randomly selects two adjacent lattice sites 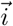 and 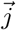, occupied by different cells 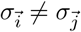. The elementary step is an attempt to copy 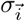 into the adjacent lattice site 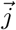, which takes place with probability

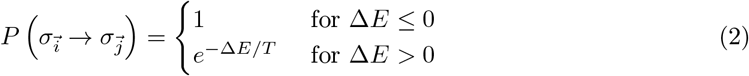

where ∆*E* is the change in functional (1) due to the elementary step considered, and the temperature parameter *T* is an arbitrary scaling factor. A Monte Carlo step of the simulation (MCS), the natural unit of time in the model, is set to *L* elementary steps – where *L* is the total number of lattice sites in the simulated area^39^. Together, Equations (1, 2) imply that cell configurations which increase the penalties in functional (1) are less likely to occur. Thus, the cell population evolves through stochastic rearrangements in accordance with the biological dynamics incorporated into the effective energy function *E*.

In the initial condition, the area of each cell was set to the equilibrium value *A*_p_, i.e., the size of each cell is 10 × 10 pixels. The total number of *N* = 3000 cells are placed in the domain: 100 *times* 30 cells in the the *x*- and *y*-dimensions, respectively. Then, the cell-free region is a 500-pixel-wide (in the *x*-dimension) empty region. This is long enough that the leading cells will not reach the end of the empty region before the end of the simulations. The surrounding wall cells are used to prevent the cells from sticking to the lattice boundaries. The barrier and wall cells (each is 10 × 10 pixels) have the CC3D "Freeze" attribute and thus, they are excluded from participating in the pixel copies of the Potts model^42^.

### 2.2 Intra-cellular dynamics

Active cell motility is powered by cytoskeletal dynamics which is regulated by cell polarity, a spatial imbalance of signaling molecules^1,17,43^. Following^35^, we represent cell polarity as a vector quantity 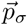, and we assume that the motile force is a nonlinear (Hill) function of polarity:

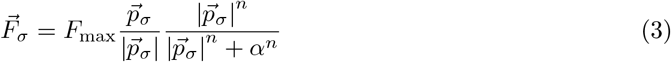

with half-saturation constant *α* > 0, and maximal motile force *F*_max_. To describe the dynamics of cell polarisation, we adopt the earlier models^24,25,35^ similar to the one recently used in^21^ as:

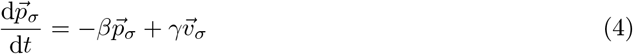

where 1/*β* is the characteristic persistence time of polarisation and the second term represents the reinforcement of polarisation through actual movement^15,44^, where 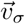 is the velocity of the center of mass of the cell *σ*. This is qualitatively equivalent to earlier models in which cell polarity aligns with cell velocity due to the inherent asymmetry created in a moving cell^25,29,45^; reviewed in^26^. Thus, the displacement of the cells is determined by the intra-cellular motile force 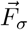 in combination with other inter-cellular interactions represented in Equations (1–4).

To mimic the initial migratory stimulus through the presence of the cell-free region to which the leading cells (i.e., the first row of cells) are exposed, when the barrier is lifted, we assign an initial polarity to the leading cells to trigger their motility. This accords with the earlier studies showing that the motile force production is initiated at the first few rows of the leading edge and travels backwards into the monolayer^19,20,46^ with a stable profile along the edge^47^.

The magnitude of the initial polarity of the leading cells is set as the steady-state polarity of a migrating single cell; see Fig. 1(a). Due to the strong nonlinearity of Equation (3), in the model single cell motility exhibits a bistable behaviour as shown in^35^; see also in Fig. 1(a). Thus, when a cell is initialized with polarity greater than the threshold parameter *α*, it will tend to move with a stable velocity and polarity. Conversely, if the initial polarity is lower than *α* the cell gradually looses its polarity and motility; see Fig. 1(a). Such bistable behavior has been experimentally observed in^48,49^. Thus, following the experiments^19,20,46,47^, at the onset of migration, when the barrier is removed, we set the polarity of the leading cells equal to the steady state value; see Table 1. We set the maximum motile force *F*_max_ = 1000, at which according to the single cell simulations the cell velocity starts to saturate; see Fig. 1(b-d) and Table 1.

**Figure 1:**
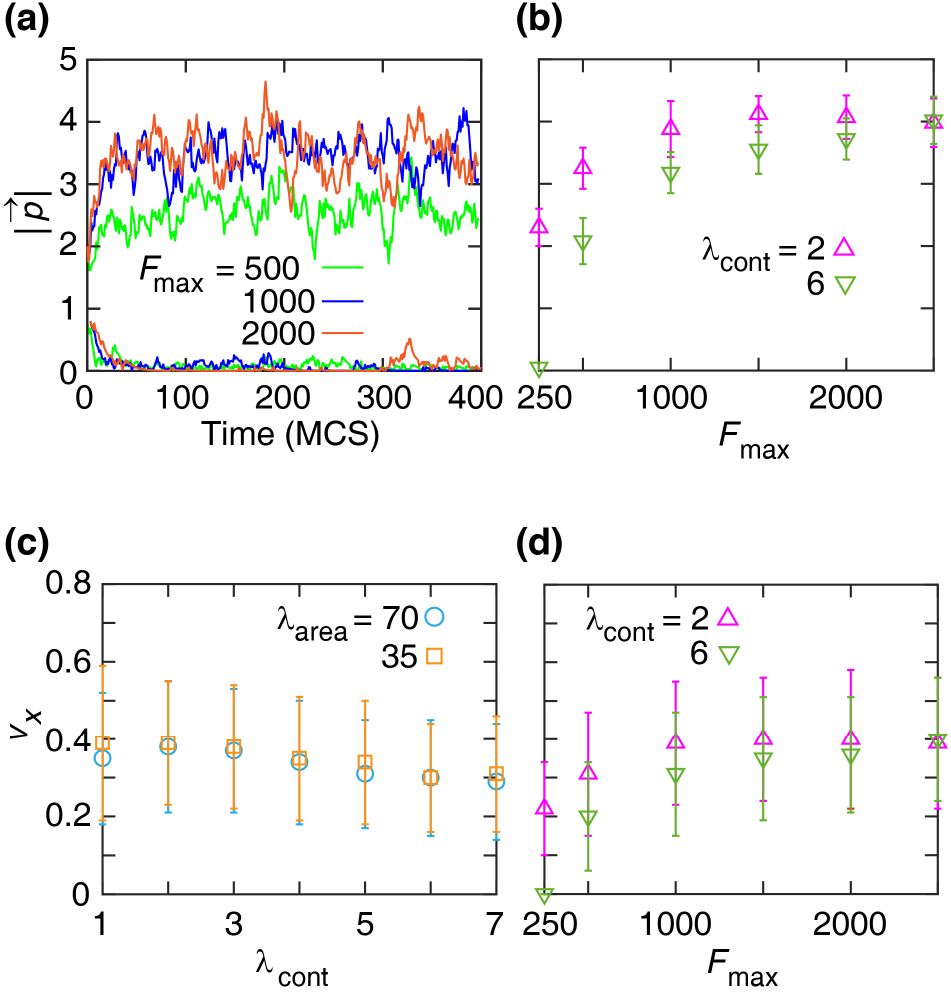
Single-cell dynamics. (a) The magnitude of polarity 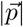 versus time plots indicate bistability. For various values of the maximal motile force *F*_max_ (see the color key), the model cell exhibits either a highly polarized or a depolarized phenotype – depending on the initial polarity value (2 or 0.8, respectively). The cortical contractility coefficient is *λ*_cont_ = 4 and the area compressibility coefficient is *λ*_area_ = 70. (b) Steady state cell polarity 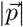 is a non-linear increasing a function of the maximal motile force *F*_max_. The steady state polarity saturates for *F*_max_ *>* 1000. The area compressibility coefficient is *λ*_area_ = 70, the contractility parameter is color-coded according to the key. see Movie 1. (c, d) Velocity *v* (*x*-component) versus *λ*_cont_, where *F*_max_ = 1000, (c); and *F*_max_, where *λ*_area_ = 70 (d). Error bars indicate standard deviation (SD). Other simulation parameters are in Table 1.

We ran each simulation in three stages. The first stage yields equilibrium cell shapes, starting from a uniform square grid initial configuration. The duration of this step is 2900 MCS, and the motile force is switched off *F*_max_ = 0. During this stage cell shapes are thus determined by the competition between cell-cell adhesion and the contractility of the cell cortex (Fig. 2) as previously reported^33,40,50^. In the "hard" regime, when contractility is strong, the cells tend to minimise their perimeter and exhibit shapes that are close to hexagons. Conversely, in the "soft" regime cell-cell adhesion dominates and cells have irregular shapes with elongated boundaries.

**Figure 2:**
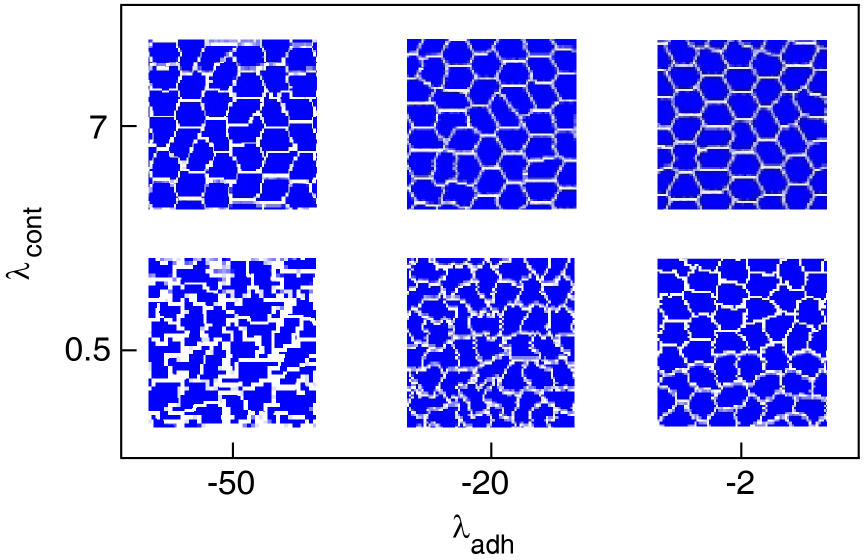
Cell shapes at a stochastic equilibrium state, for various regimes of cortex contractility *λ*_cont_ and adhesion *λ*_adh_ coefficients. Other simulation parameters are listed in Table 1.

The second stage (for 100 MCS) simulates the full dynamics of the cells, i.e. the cell polarisation coupled to the CPM through the motile force, while the cells are still confined behind the barrier. We denote the end of this stage as *t* = 0; see Fig. 3a. Finally, at the onset of stage 3 the barrier is lifted, the first row of cells are polarized (Fig. 3b, c). The simulation ends when the polarization wave traverses the domain.

**Figure 3:**
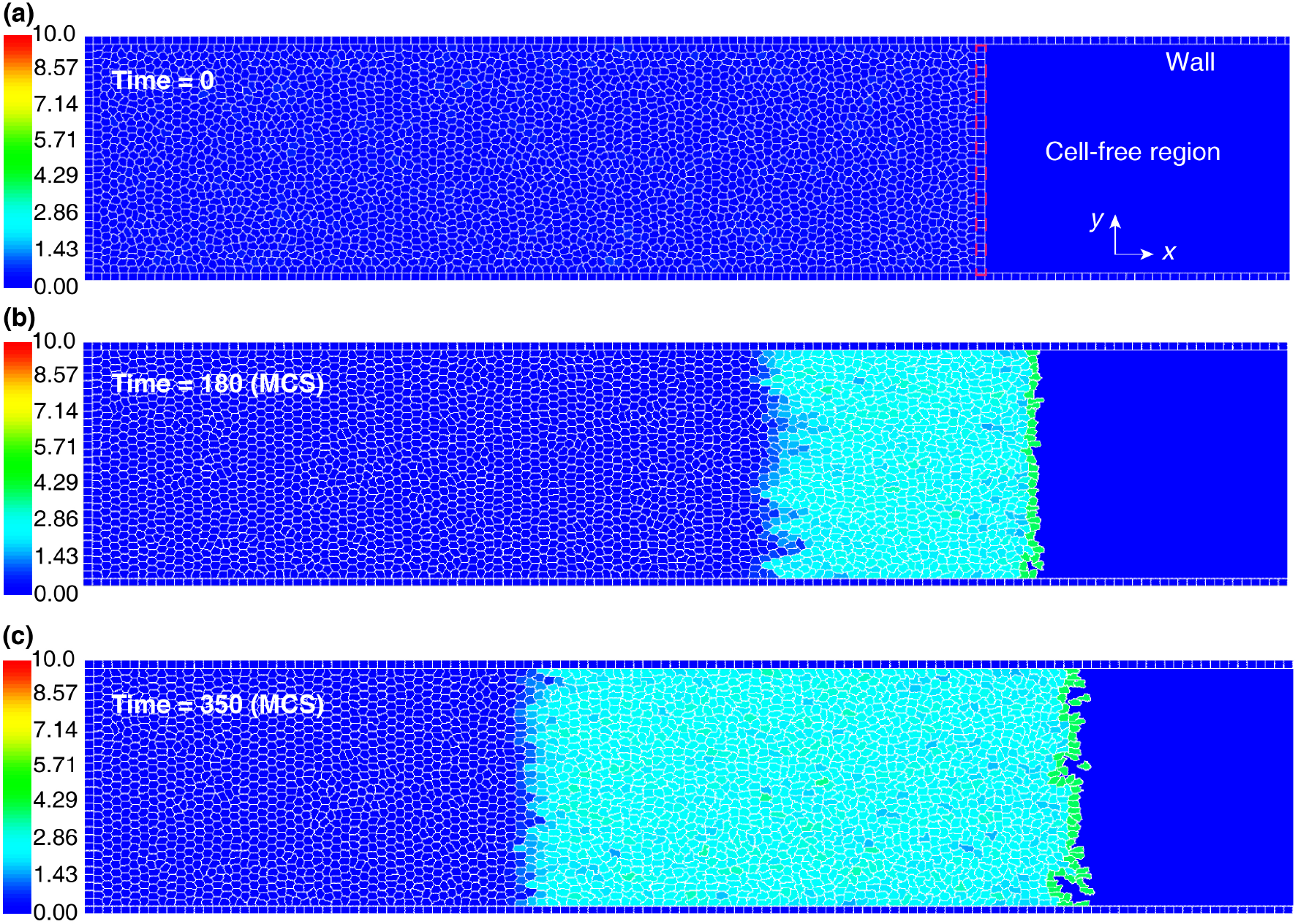
Simulation steps. (a) The barrier is lifted at *t* = 0. (b) After the barrier removal, the cells migrate into the cell-free region and cells develop polarity within an area progressively extending backward. (c) At *t* = 350 MCS, the polarization wave has propagated through half of the monolayer. Colour bar: the cell polarity magnitude 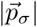.

## 3 Results and discussion

We used the model described in the previous section to systematically investigate how the various model parameters representing cellular mechanics affect collective cell movement. One interesting observation was that in a certain range of parameters, when the motile force in the CPM is switched on, cells spontaneously polarise and generate a swirling movement even in a confluent layer without a free edge (i.e. before the barrier is lifted). Typical cell trajectories corresponding to this type of characteristic swirling motion are shown in Fig. 4(a), top. The spontaneous polarisation and swirling motility happens when the contractility of the cell cortex is relatively weak and/or the polarisation threshold *α* is low. Otherwise, the cells remain unpolarised and almost stationary (Fig. 4(b), top). The corresponding phase diagram in the model parameter space is shown in Figure 5(a).

**Figure 4:**
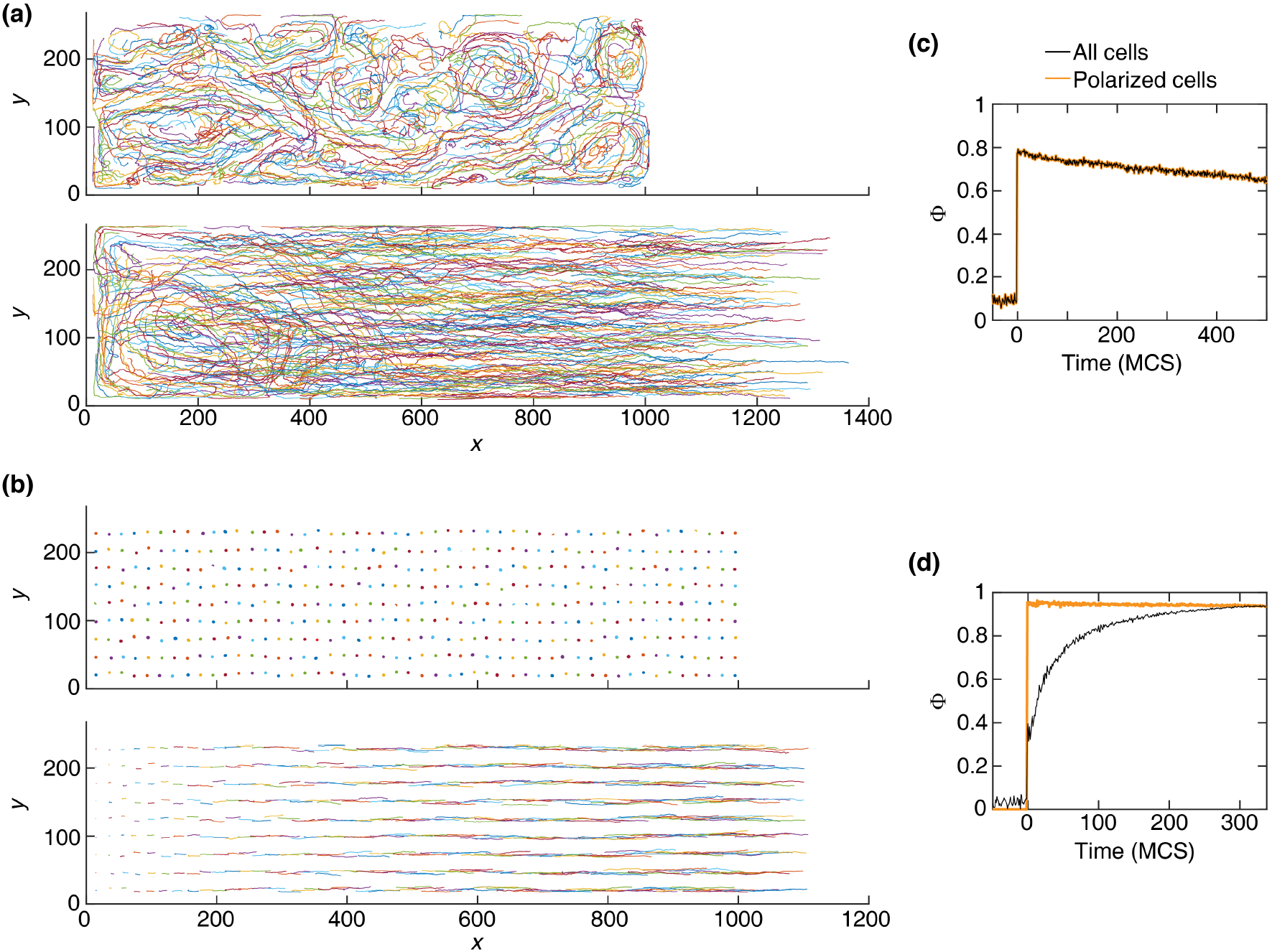
(a) Displacement trajectories of the center of mass of the cells exhibiting swirling movement, due to a weak cortex contractility *λ*_cont_ = 0.3, before (top) and after (bottom) the barrier removal; see Movie 2. Bottom: coexistence of the swirling and the migrating cells. (b) Displacement trajectories of cells with *λ*_cont_ = 3, before (top) and after (bottom) the barrier removal. Top: cells remain unpolarised. Bottom: ordered directed migration. (c) Order parameter Φ for the trajectories in (a, bottom), where the simulations are run for 500 MCS after the barrier removal. (d) Φ for the trajectories in (b, bottom), where the simulations end when the polarization wave reaches the end of the domain. *λ*_adh_ = −2. For the clarity of illustration, sample trajectories are uniformly selected from the domain. Other simulation parameters are in Table 1.

**Figure 5:**
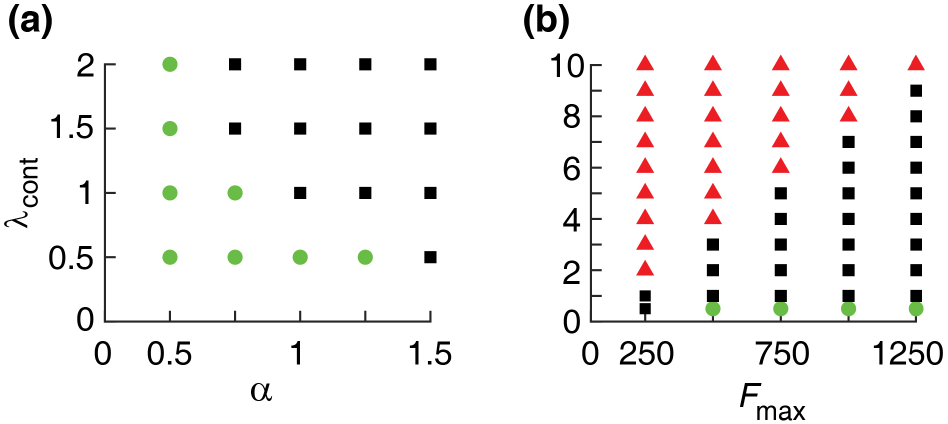
Phase diagrams. (a) Parameter region indicating the presence of spontaneous swirling motion (green circles) or stationary cells (black squares) in a confluent layer, i.e. before the barrier is lifted, when the cortex contractility coefficient *λ*_cont_ and the half-saturation constant *α* are varied. (b) Different collective cell behaviors as the contractility *λ*_cont_ and the maximal motile force *F*_max_ are varied. Swirling movements are formed at lower *λ*_cont_ ≤ 0.5 (green circles). Following the barrier removal, we observe ordered cell sheet migration (black squares), or transient cell migration when the polarization wave dies out and does not propagate into the entire monolayer (red triangles). Other simulation parameters are in Table 1.

After the barrier is lifted we observe that the cells migrate into the cell-free region; see Fig. 4(a, b), bottom. In the regime where spontaneous polarization occurs, the invasion and the swirling movements coexist – the latter is prominent in the bulk, further away from the moving edge; see Fig. 4(a), bottom. Such coexistence of the swirling and directed migration was indeed observed in experiments^51^.

To assess the effect of the swirling on the overall alignment of the migrating cells, we use an order parameter defined as:

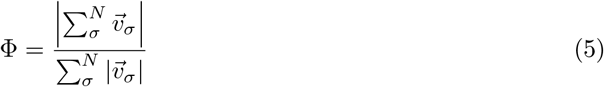

where Φ varies between 0 (uncorrelated random movement) and 1 (fully aligned migration). We find that the swirls reduce the overall alignment of the migration, captured by a decreased order parameter Φ; see Fig. 4(c). In the absence of the swirls, cell alignment gradually increases as the polarization wave propagates into the monolayer; see Fig. 4(d). Such increase in the alignment of migrating cells with the propagation of the motility wave has been observed experimentally in^20^.

Interestingly, the directed collective migration of cells into the free space requires that the motile force generated by a polarised cell (*F*_max_) is sufficiently strong. Otherwise either the polarization can not propagate to backward into the bulk or the cell layer cannot expand. In either case the migration stops after a short transient period. The minimal value of the motile force required for the propagation of the polarization wave into the bulk and for sustained directed migration increases with the cortical contractility of the cells; see Fig. 5(b).

We now focus on the parameter regime which keeps the cells stationary and unpolarised until the barrier is lifted. If the motile force is strong enough, such cells become motile and migrate into the free space. The progression of the average position of the cells is shown in Fig. 6 for various parameter values. To calculate the average position of cells, we bin the domain along the *x*-dimension, where each bin is five-cell wide, and calculate the *x*-component of the average position of cells over time. Fig. 6 shows a progression of the leading edge with constant speed, in agreement with experiments^51^. The stationary cells are recruited into the collective migration at a constant rate by the propagating polarisation wave – as predicted by our previous one-dimensional (1D) model^35^. The slope of the dashed line shows the propagation speed of the polarization wave, which decreases when the cell contractility is increased; compare the slopes of the dashed lines in Fig. 6(a, b).

**Figure 6:**
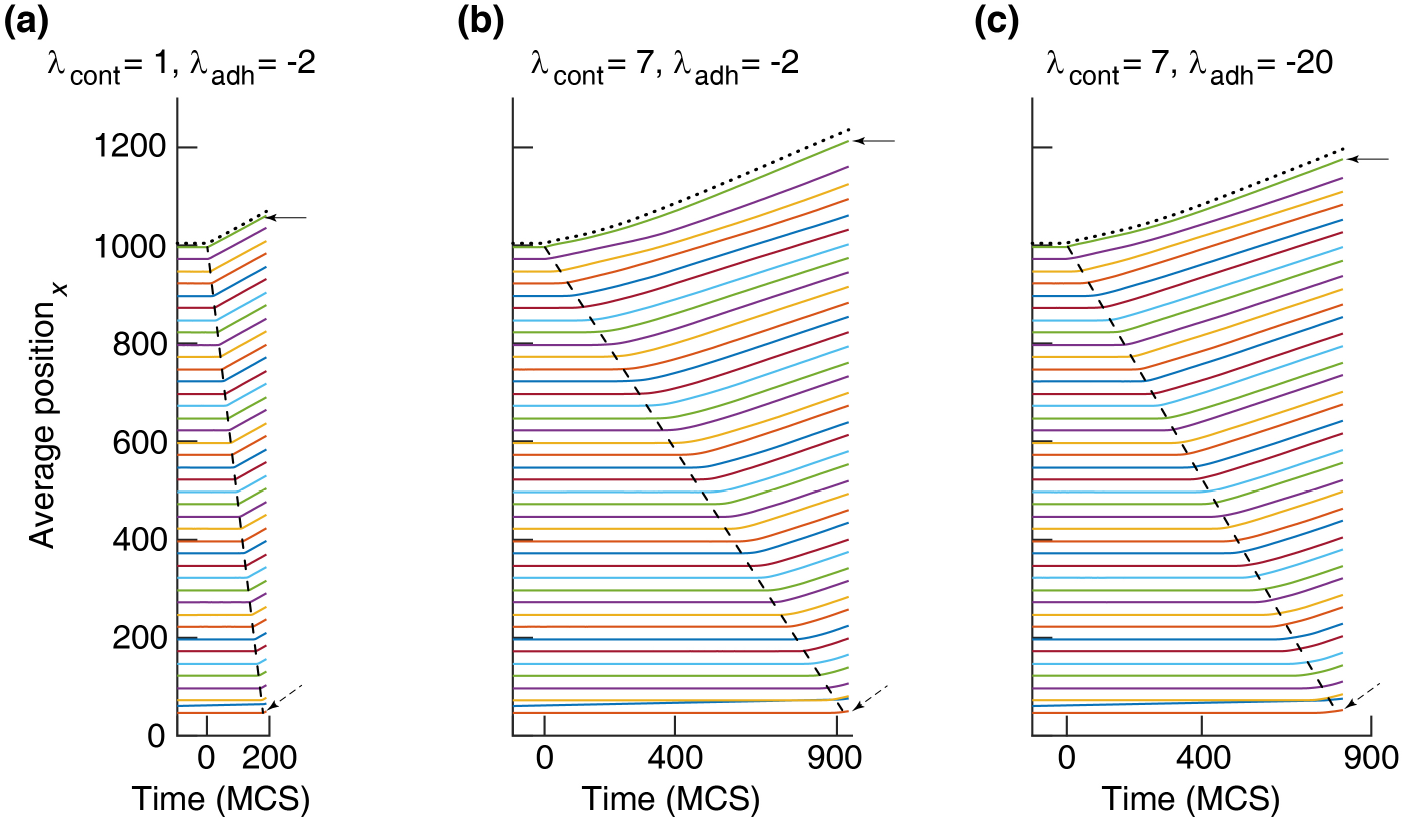
Average displacement of the center of mass of cells (*x*-component) versus time at various cortex contractility *λ*_cont_ and cell-cell adhesion *λ*_adh_. For the clarity, trajectories are averaged over five-cell wide bins along the *x*-dimension of the domain. Dotted trajectory: displacement of the leading cells only, i.e., the first row of cells from the cell-free region. Solid arrow: average position of the front bin. Dashed arrow: average position of the last bin. Dashed line indicates the propagation of the polarization as a wave, and its slope is the wave speed. *λ*_adh_: −2 (less adhesive) and −20 (highly adhesive). See Movies 3 and 4. Other simulation parameters are in Table 1.

In order to investigate how the speed of the polarization wave and the expansion velocity of the edge cells change as a function of model parameters, we ran a series of simulations and the results are summarised in Fig. 7. The polarization wave speed and the velocity of the leading edge are most strongly affected by the cortex contractility. Increasing the cortex contractility decelerates the propagation of the polarization wave and also the velocity of the leading edge. Parameters representing cell-cell adhesion and compressibility have much weaker effects (Fig. 7).

**Figure 7:**
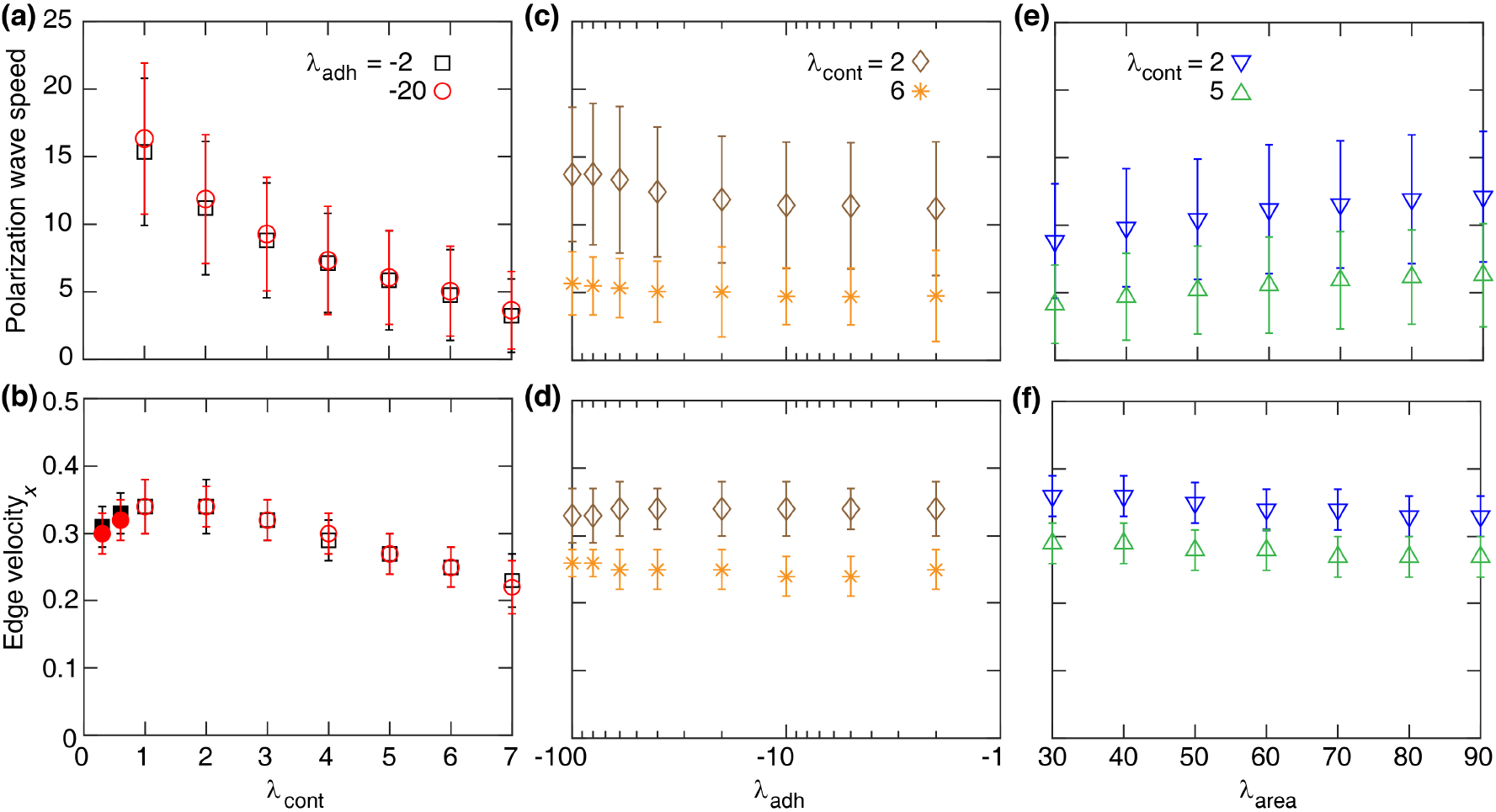
The polarization wave speed (top row) and the *x*-component of the edge velocity (bottom row) versus the cortex contractility coefficient *λ*_cont_ (a, b), the cell-cell adhesion coefficient *λ*_adh_ (c, d), and the area strain *λ*_area_ (e, f). Each symbol is derived from a single simulation run and corresponds to mean ± SD. In (b), the filled symbols are at *λ*_cont_ = 0.3 and 0.6 where the swirling movements are formed. Other simulation parameters are in Table 1

At low cortex contractility the edge velocity (~ 0.3) is not sensitive to the presence or absence of swirling motion (Fig. 7(b), filled symbols). This finding is consistent with the experimental observation that the average velocity of the migrating cell front did not show much variation in the presence of swirls^51^. Consistent with earlier reports^52^, Fig. 7(c-f) also indicates that cell-cell adhesion and the area compressibility have little effect on the polarization wave speed and the edge velocity.

For certain parameter values the simulations indicate patterning instabilities at the free edge: some of the leading cells may develop multicellular migration fingers, which was also observed experimentally in^41^. We find that migration fingers develop when the cortex contractility is strong. When the cortex contractility is weak (e.g., *λ*_cont_ = 2), the energy cost for the advancement of the leading cells is low, resulting in a smooth migration of the leading edge of the monolayer; see Fig. 8(a). However, with strengthening the cortex contractility (e.g., *λ*_cont_ ≈ 6 or larger), the displacement of the leading cells is somewhat restricted, the stochastic fluctuations are amplified resulting in the development of migration fingers; see Fig. 8(b, c). This agrees with the experimental observations^41^ that the tension in the cortical actomyosin ring prevents the initiation of new leader cells, and the limited number of leader cells results in the development of the migration fingers.

**Figure 8:**
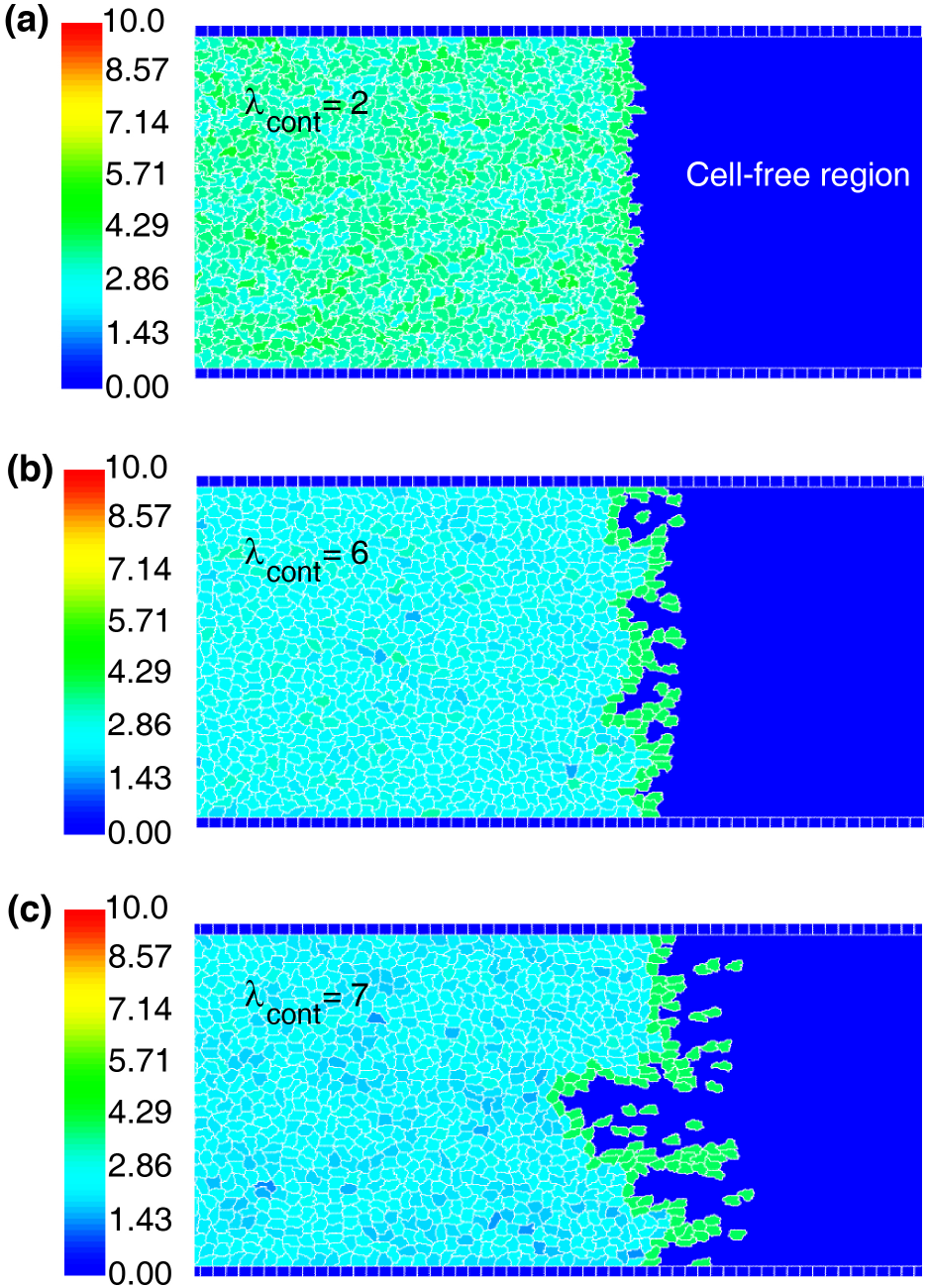
Leading edge migratory dynamics. The leading edge destabilizes into the migration fingers, when the cell cortex contractility *λ*_cont_ increases from 2 (a) to 6 (b) and then to 7 (c). Screenshots are taken at the end of the simulations. *λ*_adh_ = *−*20. Other simulation parameters are in Table 1. Colour bar: the cell polarity magnitude 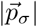.

## 4 Summary and outlook

In this paper, we studied the collective migration of epithelial cells in a confluent monolayer exposed to free space. Our simulations identified that the cortex contractility, not adhesion of the cells is a key model parameter that controls the transitions between swirling movements and well aligned cell sheet migration. Strong contractility can promote the formation of migration fingers, block the propagation of the polarization wave, and inhibit the collective migration.

Our model also indicates the role of the cells cortex contractility in regulating the formation of the swirls and further predicts that the swirls are more likely to form when the cortex contractility is weak; see Fig. 4(a, b). This is explained using Equations (1–4). Cells with a weak cortex contractility (e.g., *λ*_cont_ = 0.3) can more easily change their shape which can also lead to the displacement of the center of mass of the cells. This displacement can spontaneously polarize the cell, due to the bistable behaviour of the cell polarity 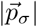; see Fig. 1(a). The spontaneous cell polarization becomes more likely when the polarization threshold *α* is lower, resembling the experimental observations that the intensity of the polarization promotes the appearance of the swirls^53^. Then, the motile force 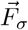 generated by a spontaneous polarization is transmitted to the neighbouring cells and form swirling movements. This suggests that sufficiently strengthening the cells cortical contractility would increase the overall alignment of the migration by transitioning the swirling movements into a directed cell sheet migration; see Fig. 4(a, b).

On the other hand, excessive strengthening of the cells cortex contractility can block the collective migration in a monolayer; see Fig. 5(b) and Movie 5. A tight cortical contractility results in symmetric hexagon-like cell shapes corresponding to the "hard" regime (Fig. 2) that blocks the cells polarization by limiting their deformation and motility. The "hard" cell shapes have relatively little displacement of the center of mass of the cells, corresponding to a weak cell polarity and motile force. This blockage of the active motility can also be caused by increasing the decay rate of the cell polarization *β*. At high *β*, the cells do not possess non-zero steady state for the polarity and their polarity magnitude can only stabilise at zero; see Fig. 1(a) and our discussion in^35^. Then, starting with an initial polarity, the cells gradually loose their polarity and stop moving. Therefore, the polarization wave becomes unstable and disappears gradually after the barrier is lifted; see Movie 6.

In the present study, we focused on the migration of cells in the absence of cell proliferation, and where all the cells had the same constant mechanical characteristics. Future works may consider the effects of the cell proliferation combined with active cell migration, on the collective movement and the propagation of the polarization wave. It is also interesting to consider the adaptive responses of cells to the environment, where the cellular properties can vary. For instance, the alignment of the cells migration may be enhanced by coupling the cortical contractility to the polarization threshold in order to inhibit the swirling movements; see Fig. 5(a, b). Likewise, recent studies on the efficiency of the wound healing have shown that a monolayer can effectively cover the wound region and achieve a permanent gap closure when the formation of finger-like shapes is supressed^54^. Accordingly, a coupling of the cortical contractility to the morphology of the leading edge in order to maintain a moderate cortical contractility could be studied in the context of a wound closure assay.

## Acknowledgement

This work was initiated during the workshop "Virtual tissues, progress and challenges in multicellular systems biology" organised by J. Osborne. We acknowledge very useful discussions with J. Glazier on the implementation of our model in CompuCell3D.

## Author contributions

ZN conceived the idea. HK carried out the model development, analysis, and prepared the first version of the manuscript. All the authors, HK, AC, ZN, contributed to the interpretation of results and improvement of the manuscript.

## Supporting Information

**Movie 1**. Single-cell movement, with cortex contractility *λ*_cont_ = 2 and maximal motile force *F*_max_ = 1000. Other simulation parameters are in Table 1.

**Movie 2**. Cells exhibit swirling motions, due to low contractility *λ*_cont_ = 0.3. Colour bar: polarity magnitude 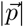. The first simulation stage is illustrated as “Traction force is off”. The note “Traction force is on” corresponds to the second and third simulations stages. Other simulation parameters are in Table 1.

**Movie 3**. Collective cell migration, where cells cortical contractility *λ*_cont_ = 1 and *λ*_adh_ = −2. Other simulation parameters are in Table 1.

**Movie 4**. Collective cell migration, where cells cortical contractility *λ*_cont_ = 7 and *λ*_adh_ = −2. Other simulation parameters are in Table 1.

**Movie 5**. Depolarization of the monolayer, due to high cell contractility *λ*_cont_ = 8. *λ*_adh_ = −10. Other simulation parameters are in Table 1.

**Movie 6**. Depolarization of the monolayer, due to high depolarization rate *β* = 0.3. *λ*_cont_ = 1, *λ*_adh_ = *−*10, and *λ*_area_ = 30. Other simulation parameters are in Table 1.

